# Compressive forces stabilise microtubules in living cells

**DOI:** 10.1101/2022.02.07.479347

**Authors:** Yuhui Li, Ondřej Kučera, Damien Cuvelier, David M. Rutkowski, Mathieu Deygas, Dipti Rai, Tonja Pavlovič, Filipe Nunes Vicente, Matthieu Piel, Gregory Giannone, Dimitrios Vavylonis, Anna Akhmanova, Laurent Blanchoin, Manuel Théry

**Affiliations:** Univ. Paris, INSERM, CEA, UMRS1160, Institut de Recherche Saint Louis, CytoMorpho Lab, Hôpital Saint Louis, 75010 Paris, France; Univ. Grenoble-Alpes, CEA, CNRS, INRA, Interdisciplinary Research Institute of Grenoble, Laboratoire de Phyiologie Cellulaire & Végétale, CytoMorpho Lab, 38054 Grenoble, France; Institut Curie, UMR144, Paris, France; Institut Pierre-Gilles de Gennes, Paris, France; Department of Physics, Lehigh University, Bethlehem PA 18015, USA; Cell Biology, Neurobiology and Biophysics, Department of Biology, Faculty of Science, Utrecht University, Utrecht, The Netherlands; Institut Interdisciplinaire des Neurosciences, CNRS UMR 5297, Université de Bordeaux, F-33000 Bordeaux, France

## Abstract

Cell mechano-sensation and adaptation are supported by the actin network. The microtubule network is not considered to be directly sensitive to mechanical forces acting on a cell. However, recent studies on isolated microtubules *in vitro* have shown that bending forces have an impact on their structure, composition and lifespan, suggesting that, in a cellular context, microtubules may react to mechanical forces. We tested this hypothesis in living cells by subjecting them to cycles of compressive forces and found that microtubules became distorted, less dynamic and more stable. This mechano-stabilisation depends on CLASP2, which relocates from the end to the deformed shaft of microtubules. These results demonstrate that microtubules in living cells have mechano-responsive properties that allow them to resist and even counteract the forces to which they are subjected.

---

Mechanical forces not only deform cells but also instruct them and, thereby, regulate the functional organisation of tissues (*1*). In particular, mechanical forces regulate cell migration (*2*), cell cycle progression (*3*), mitotic spindle orientation (*4*) and stem cell differentiation (*5*). In various organisms, these processes are often coupled to the reorganisation of the microtubule (MT) network and, in particular, to its local stabilisation in specific subcellular regions (*6*–*9*). Whether and how such regulation is a direct consequence of mechanical forces is not known.

The mechanosensitive properties of cells and their mechano-response are thought to depend mainly on cell adhesion complexes and the actin cytoskeleton (*10*–*12*). In particular, the adhesion and actin networks are known to strengthen in response to mechanical forces (*13*). However, the MT network also changes in response to physical constraints, which modulate MTs growth rate (*14*, *15*) and direct their orientation (*16*). Mechanical forces also have an impact on the post-translational modification of tubulin (*17*, *18*), which modulates MT stiffness and deformation under load (*19*, *20*). It is not yet clear whether these changes are an indirect consequence of the interaction of microtubules with the actin network (*21*) or whether cytoplasmic microtubules have autonomous mechanosensitive properties in cells.

*In vitro* experiments on isolated MTs have shown that pressure on a growing MT tip can induce MT disassembly (15). In contrast, physical injury induced by laser ablation or bending forces can trigger self-repair of the MT lattice (*22*), which can protect MT from disassembly (*23*, *24*). These *in vitro* experiments suggest that MTs *in vivo* may have specific mechano-responsive properties, but direct evidence in living cells has been lacking, and the impact of mechanical signals on microtubule stability *in vivo* has not yet been discovered.

The aim of our study was to apply controlled mechanical forces to living cells and test the response of the MT network. We used a thin silicone sheet clamped to a motorised stage to apply stretch and compression cycles (SCC) to cells (*25*). We implemented a method to print adhesive micropatterns on the silicone sheet to normalise the response of the cells and to study the cytoskeleton response independently of resultant cell-shape change (Figure 1A and S1A-B). The shape of the membrane and its clamping in the movable elements ensured a uniaxial SCC without compensation by perpendicular compression in the central part (Figure S1 C-H). We first tested the resistance of the cell to the deformation by applying SCC of different magnitudes (10 to 60%). The cells detached at 40%, but there was no visible impact on the shape of the cell at 10% strain (Supplementary Figure S2 A, B). We also tested the effect of SCC frequency on cell detachment. The cells started to detach at 0.2 Hz but held up well to 10% strain cycles at 0.1 Hz (Supplementary Figure S2 C, D). We, therefore, chose to constrain the cells by applying 10 cycles of 10% SCC for 2 minutes (i.e. 0.08 Hz).

**Figure 1.**
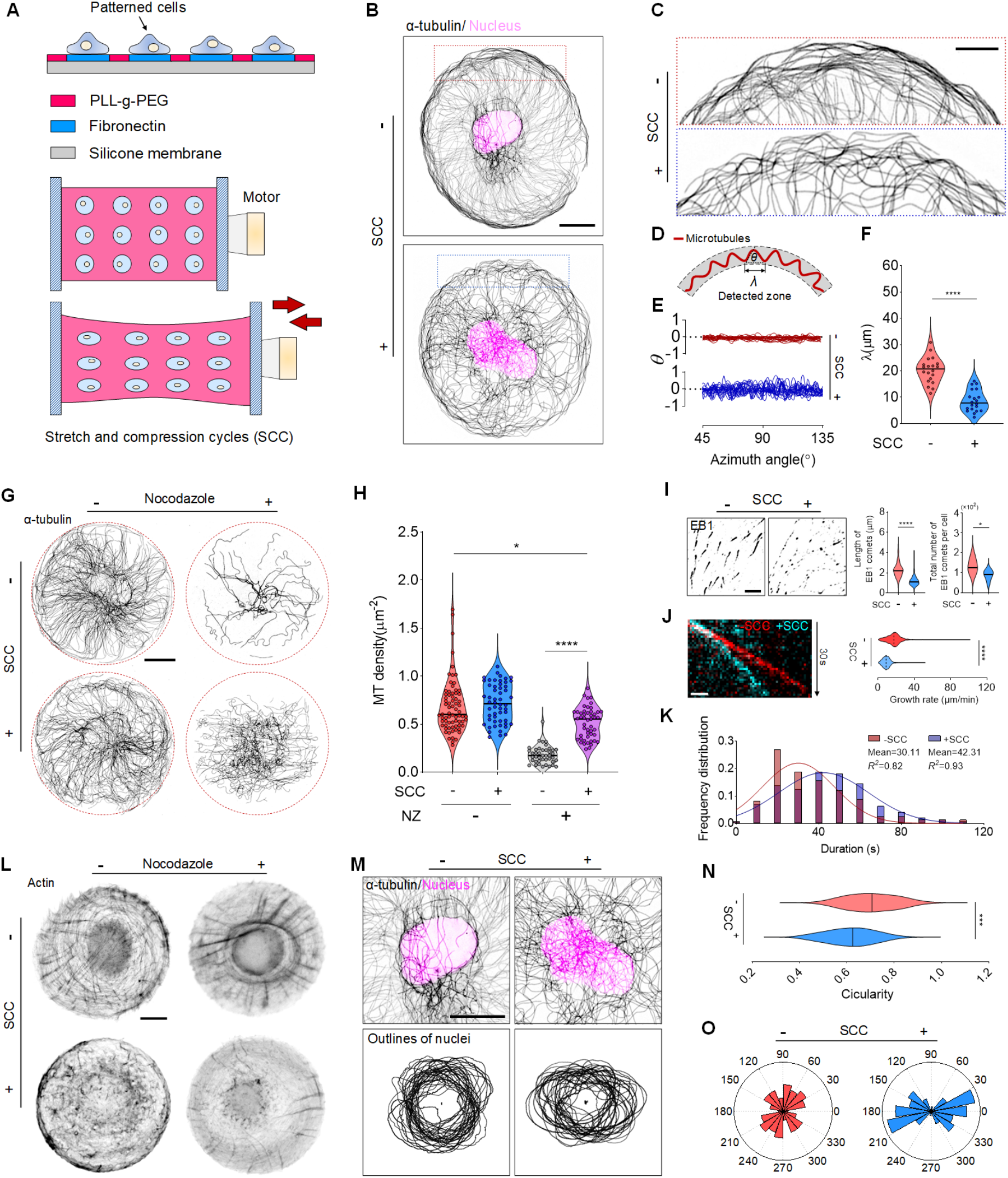
Microtubules are deformed and stabilised in response to stretch and compression cycles (SCC) in RPE1 cells. (A) Illustration of the SCC assay. RPE1 cells spread on a PMDS membrane micropatterned with disk-shaped fibronectin (50 μm in diameter). A uniaxial stretching device comprises a holding arm and stretching arm connected to a mechanical motor. (B) Inverted immunofluorescence (IF) images of individual micropatterned RPE1 cells without/with 10 cycles of SCC at 10% strain, stained for α-tubulin (grey) and DNA (magenta). (C) Zoomed-in images of the microtubules in peripheral regions of representative cells in panel B. (D) Schematics illustrating the characteristic wavelength quantification of deformed microtubules. (E) Azimuth orientation graph of microtubules representing the MT waviness in cortical regions of 20 cells without/with SCC (three independent experiments). (F) Violin plots of MT characteristic wavelength in two conditions; -SCC/+SCC: without/with SCC (n = 20 cells, three independent experiments). Asterisks indicate Mann-Whitney test significance values; \*\*\*\**p* < 10^−4^. (G) Inverted IF images of RPE1 cells stained for α-tubulin in four conditions. - SCC/+SCC: without/with SCC, −NZ/+NZ: without/with nocodazole treatment. The SCC axis is parallel to 0°. (H) Violin plots of microtubule density of RPE1 cells in varying conditions (−SCC/−NZ: n =72 cells, +SCC/−NZ: n =54 cells, −SCC/+NZ: n =42 cells, +SCC/+NZ: n =49 cells, three independent experiments). Asterisks indicate significance values from the Kruskal-Wallis test with Dunn’s multiple comparison test; *\*\*\*\*p* < 10^−4^, \**p* < 0.05. (I) Representative inverted IF images of cells (left) without/with SCC, stained for EB1 and violin graph of the length (middle) (n = 150 comets from 11 cells, three independent experiments) and total number (right) (n = 7 cells, three independent experiments) of EB1 comets. Asterisks indicate Mann-Whitney test significance values; \*\*\*\**p* < 10^−4^, \**p* < 0.05. (J) Representative kymographs (left) and growth rates quantification (right) of GFP-EB1 in cells without/with SCC. (n > 500 comets tracked from 7 cells, two independent experiments). Asterisks indicate Mann-Whitney test significance values; \*\*\*\**p* < 10^−4^. (K) Histogram of EB1 lifetime distribution with Gaussian fit functions in cells without/with SCC. (n = 150 comets from 11 cells, three independent experiments). (L) Inverted fluorescence images of cells in varying conditions, stained for actin. (M) Zoomed-in images of central regions of representative cells in panel B (top) and projected images of nuclear outlines (bottom) in cells with/without SCC (+SCC: n =31 nuclei, –SCC: n =36 nuclei, three independent experiments). (N) Violin graph of nuclear circularity in cells without/with SCC (–SCC: n =48 nuclei, +SCC: n =37 nuclei, three independent experiments). Asterisks indicate Mann-Whitney test significance values; \*\*\**p* < 10^−3^. (O) Polar graph of nucleus orientation in cells without/with SCC. (n =39 nuclei, three independent experiments). Scale bar, 10 μm (B, G, L and M), 5 μm (C) or 2 μm (I and J). Medians were depicted in violin plots (solid line).

The MT network resisted the SCC and remained globally isotropic despite a few stereotypical changes (Figure 1B, see more examples in Supplementary Figure S2 E, F). In particular, we noticed the appearance of curved MTs along the cell periphery (Figure 1C). Indeed, measurement of the local orientation of the MTs and the characteristic wavelength (Figure 1D) revealed an increase in orientation variations (Figure 1E) and a reduction in the characteristic wavelength (Figure 1F) in sections along the SCC axis. These results suggest that the limited longitudinal elasticity of MTs does not allow them to expand during stretching phases, but that friction with the surrounding cytoplasm promotes their bending with a short wavelength during compression phases (*26*). As previous reports have shown an interesting correlation between curvature, stiffness and stability of microtubules (*27*–*29*), we next tested whether SCC has an impact on MT stability.

Nocodazole (NZ) is a cell-permeable drug that binds to tubulin dimers and thus triggers the disassembly of dynamic microtubules (*30*). We, therefore, measured the stability of MTs by testing their resistance to low doses of NZ and compared unstressed cells to those subjected to SCC (Figure 1G). Interestingly, SCC had no impact on the overall density of MT (Figure 1H, - NZ). However, it drastically increased the amount of MTs that could withstand NZ (Figure 1H, +NZ). To further characterise MT dynamics, we analysed the binding of EB1 to the plus ends of MTs since its local accumulation is a faithful indicator of MT growth (*31*) (Figure 1I). EB1-positive comets at the plus ends of MTs appeared shorter and less numerous after SCC (Figure 1I, Supplementary Figure S3A). Interestingly, additional EB1-positive spots appeared along the MT shaft, at some distance from the end (Supplementary Figure S3B). In addition, the use of a thin stretchable silicone layer that can slide over a glass surface allowed live imaging after SCC (*32*). It revealed that EB1 comets moved more slowly (Figure 1J) and had a longer lifetime (Figure 1K), indicating that MT growth was reduced. Both sets of data showed that SCC had a strong stabilising effect on MTs.

It should be noted that SCC has a well-characterised effect on the reorientation of actin filament bundles along the SCC axis at short time scales (several minutes) and along the minimum strain axis at longer time scales (several hours) in response to a wide range of frequencies (mHz to Hz) (*25*, *33*, *34*). As the actin network also influences MT stability (*17*, *35*), the stabilisation of MTs along the SCC axis that we observed might be due to a reorganisation of the actin network. However, at this frequency and strain, actin bundles did not reorient. On the contrary, they were massively destroyed by SCC (Figure 1L, bottom left). Furthermore, the remaining actin structures in the SCC-treated cells did not form additional bundles in response to NZ as they did in SCC-untreated cells (Figure 1L, bottom right). The possible role of actin network destruction in response to SCC was further assessed by treating the cells with Cytochalasin D to induce actin network disassembly. This treatment did not increase the amount of nocodazole-resistant MTs (Supplementary Figure S4), confirming that MT stabilisation was not a consequence of actin reorientation or destruction by SCC.

The interaction of MTs with the nucleus could also be involved in their stabilisation. Indeed, the nucleus is a major load-bearing structure in cells (*36*). It interacts with MTs (*37*), and its deformation is associated with the translocation of biochemical signals from the cytoplasm (*38*). Its deformation by SCC may therefore have an indirect impact on MT stability. Therefore, we tested whether the shape of the nucleus was affected in our SCC range (Figure 1M). We found that nuclei appeared elongated along the SCC axis (Figure 1M-O), suggesting that they undergo a non-elastic deformation in response to SCC. To challenge this hypothesis of an indirect role of a nucleus and to further assess the specific response of MTs to SCC, we enucleated the cells by centrifugation and plated the resulting cytoplasts onto micropatterns on stretchable membranes (*39*).

Cytoplasts appear to be more resistant to SCC than cells (Supplementary Figure S5A). However, strain larger than 20% seems to induce some fragmentation of the MT network (Supplementary Figure S5B), so we decided to stimulate the cytoplasts to the same level as cells, i.e. 10%. Interestingly, the reorientation and deformation of MTs by SCC were more pronounced in cytoplasts than in cells (Figure 2). The entire network architecture was impacted (Supplementary Figure S6). We analysed the orientation of MT with respect to the cytoplast radius (Figure 2A). The MTs have reoriented from a rather radial to a circular organisation (Figure 2B, C). In addition, the sections of these circular MTs that were aligned with the SCC axis appeared more curved (Figure 2D, E). Furthermore, the proportion of NZ-resistant microtubules was significantly increased by mechanical SCC (Figure 2F, G, Supplementary Figure S6). The average density of NZ-resistant MTs was about five times higher in cytoplasts subjected to SCC than in untreated cytoplasts. As in the case of cells, mechanically-induced stable MTs tended to align with the SCC axis (Figure 2F, H). These results show that the stabilisation of MTs by SCC was independent of nucleus deformation and likely a specific response of MTs to external stretch-compression forces.

**Figure 2.**
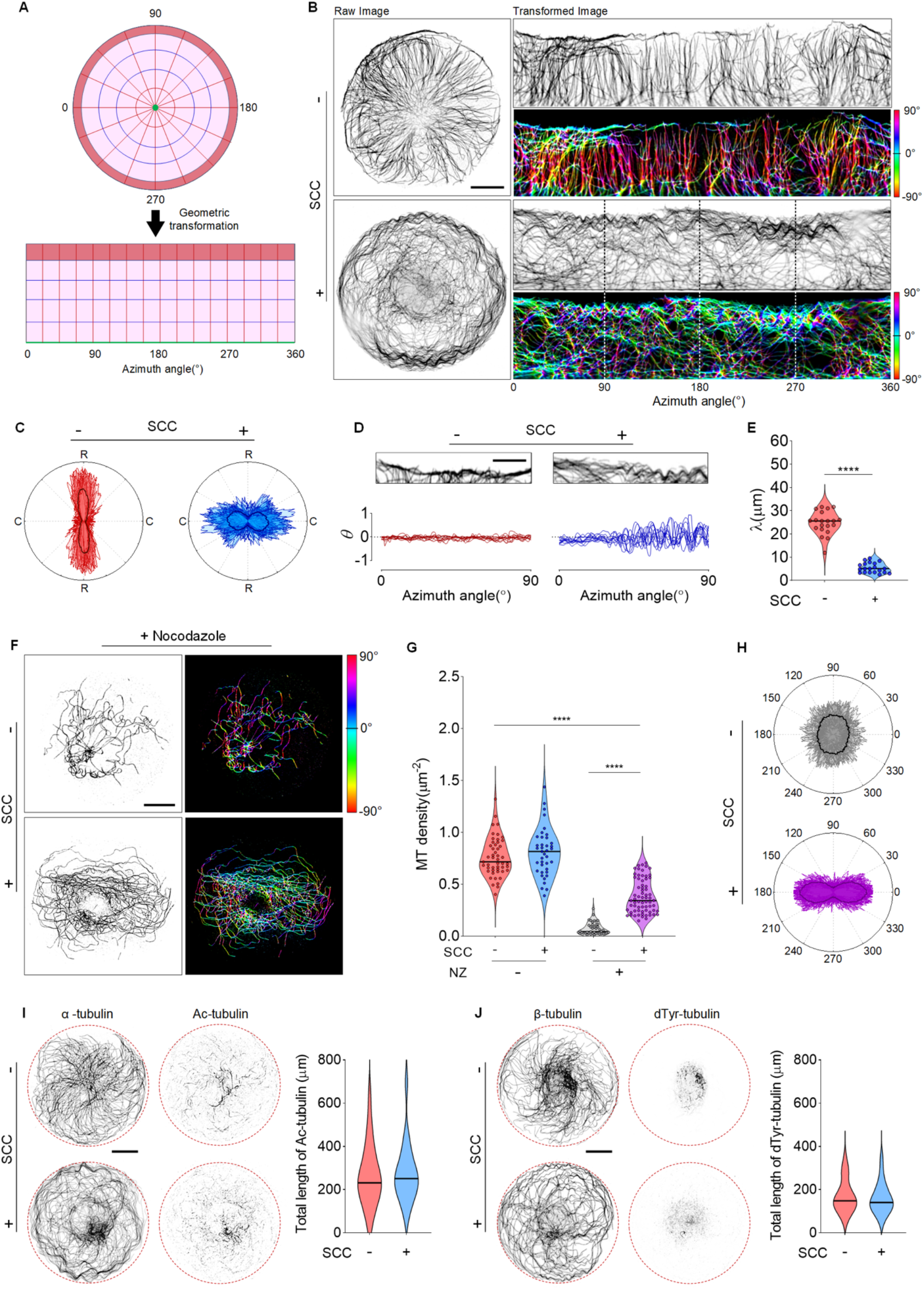
Microtubules are stabilised in response to SCC in enucleated RPE1 cells (cytoplasts). (**A**) Illustration of geometrical transformation from a disk to rectangle-shape. (B) Inverted IF images of RPE1 cytoplasts without/with SCC, stained for α-tubulin (left) and corresponding transformed images (right). The colour-coded images indicate the local microtubule orientation. The arrows in IF and colour-coded images indicate the corresponding SCC axis in both raw and transformed images. (C) Orientation polar graph of the microtubules in cytoplasts in transformed images without/with SCC (n =31 cytoplasts, four independent experiments). R and C represent a radial and circumferential orientation, respectively. Means are depicted. (D) Inverted IF images of cortical regions (azimuth angle are from 0° to 90°) of RPE1 cytoplasts without/with SCC, stained for α-tubulin (top) and corresponding azimuth orientation analysis of microtubules (bottom) that indicate the local MT waviness (n =11 cytoplasts, four independent experiments). (E) Violin plots of MT characteristic wavelength in two conditions. (n =11 cytoplasts, four independent experiments). Asterisks indicate Mann-Whitney test significance values; \*\*\*\**p* < 10^−4^. (F) Inverted IF images of RPE1 cytoplasts stained for α-tubulin (left) and corresponding colour-coded orientation images in −SCC/+NZ and +SCC/+NZ conditions. (G) Violin plots of microtubule density of RPE1 cytoplasts in varying conditions (−SCC/−NZ: n =49 cytoplasts, +SCC/−NZ: n =39 cytoplasts, −SCC/+NZ: n =42 cytoplasts, +SCC/+NZ: n =68 cytoplasts, three independent experiments). Asterisks indicate significance values from the Kruskal-Wallis test with Dunn’s multiple comparison test; \*\*\*\**p* < 10^−4^. (H) Orientation polar graph of microtubules in RPE1 cytoplasts in – SCC/+NZ and +SCC/+NZ conditions (n =20 cytoplasts, three independent experiments). Means are depicted. (I) Inverted IF images of RPE1 cytoplasts without/with SCC, stained for α-tubulin and acetylated-tubulin (left) and corresponding violin plot of total microtubule length quantification (right, n =20 cytoplasts, three independent experiments). Significance values are obtained from the Mann-Whitney test; *p* = 0.7127. (J) Inverted IF images of RPE1 cytoplasts with/without SCC, stained for β-tubulin and dTyr-tubulin (left) and corresponding violin plot of total microtubule length in RPE1 cytoplasts by using SOAX segmentation (right, n =20 cytoplasts, three independent experiments). Significance values are obtained from the Mann-Whitney test; *p* = 0.6734. Scale bar, 10 μm (B, F, I and J) and 5 μm in D. Medians were depicted in violin plots (solid line).

Microtubule stability is often correlated with post-translational modifications of tubulins (*28*, *40*). We, therefore, wondered whether the mechanical stabilisation we observed was associated with post-translational modifications. We measured the amount of glutamylated and tyrosinated tubulin (Figure 2I) as well as the amount of acetylated tubulin (Figure 2J) and found no effect of SCC. This observation showed that stabilisation was not a consequence of post-translational modifications of tubulins but did not exclude that a longer SCC could have induced such modifications, as occurs during cardiomyocytes maturation (*41*). Our results also suggested that the stabilisation was mediated by another pathway. In particular, the tortuous shape of NZ-resistant MTs suggests that their stabilisation could arise from local deformation rather than end-stabilisation. Furthermore, the re-localisation of EB1 along the length of MTs in cells subjected to SCC suggests that stabilisation may be associated with the protective role of +TIP complexes acting on the lattice of the MTs (*42*, *43*).

We investigated this possibility and analysed the spatial distribution of EB1 in cytoplasts (Figure 3A). Consistent with the experiments in cells, EB1 was localised to the growing ends of MTs under control conditions and along the entire length of MTs in response to SCC (Figure 3B). We also analysed the localisation of CLASP2, which has been shown to play a protective role on the damaged lattice of MTs in vitro (*43*). As with EB1, it relocated from the MT ends to the MT shaft in response to SCCs. Furthermore, the amount of CLASP2 on MTs appears to be specifically enriched along NZ-resistant MTs in cytoplasts subjected to SCC (Figure 3C). We implemented an image segmentation tool to quantify this effect (Methods). It allowed us to skeletonise MTs (*44*) and detect CLASP spots to measure the ratio between CLASP bound to MT versus cytoplasmic CLASP signal (Supplementary Figure S6). This analysis revealed clear recruitment of CLASP2 along the length of MTs in response to SCC (Figure 3D) and particularly on curved regions of MTs (Figure 3E). Furthermore, NZ-resistant MTs also showed a higher density of CLASP2 compared to the entire pool of MTs (Figure 3D), and this density was enriched by SCC (Figure 3D, Supplementary Figure S6B). Furthermore, heat maps of the entire cytoplasts revealed a striking correlation between the MT-associated CLASP2 and regions where the MT orientations were highly variable (Figure 3F). Overall, these results showed that CLASP2 was enriched on MTs that were bent by external forces and became resistant to NZ.

**Figure 3.**
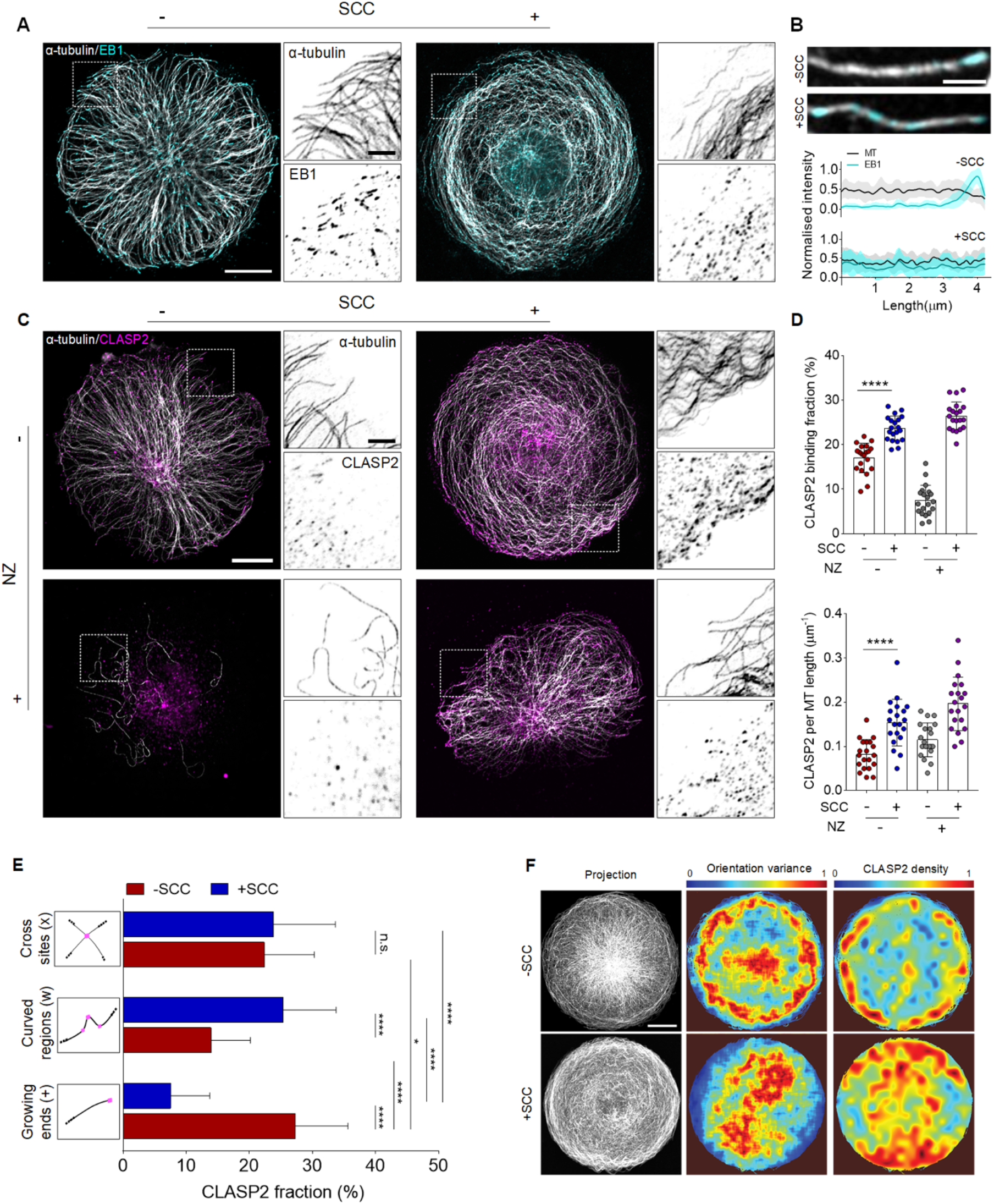
CLASP2 is enriched along microtubules in response to SCC. (A) IF images of RPE1 cytoplasts with/without SCC, stained for α-tubulin (grey) and EB1 (cyan). Dashed rectangle designates region shown with α-tubulin or EB1 channel isolated. (B) Representative IF images of individual microtubules and EB1 comets in cytoplasts with/without SCC (top) and corresponding line-scan profiles of normalised fluorescence intensity (bottom) (n =20 individual microtubules from 10 cytoplasts, three independent experiments). Means are depicted. (C) IF images of RPE1 cytoplasts with/without SCC, stained for α-tubulin (grey) and CLASP2 (magenta). Dashed rectangle designates region shown with α-tubulin or CLASP2 channel isolated. (D) Scatter plot of binding fraction and the number of CLASP2 along the length of microtubules in four conditions (n =20 cytoplasts, three independent experiments). Data are shown as mean ± s.d. Asterisks indicate significance values from the Mann-Whitney test; \*\*\*\**p* <10^−4^. (E) Histogram of the fraction of CLASP2 binding on specific regions including growing ends, curvature peak and crossing sites of microtubules (schematics illustrating the specific binding regions) (n=40 ROI regions of 10 cytoplasts, three independent experiments). Data are shown as mean ± s.d. Asterisks indicate significance values from the Kruskal-Wallis test with Dunn’s multiple comparison test; \*\*\*\**p* < 10^−4^, \**p* < 0.05, for *n.s.* (not significant) *p* > 0.1. (F) Projected IF images of RPE1 cytoplasts without/with SCC, stained for α-tubulin (left), corresponding heatmap of microtubule orientation variance (middle) and the density of microtubule-associated CLASP2 (n=15 cytoplasts, three independent experiments). Scale bar, 10 μm (A, C and F), 2 μm (zoomed-in of A and C) or 1 μm (B).

CLASP2 is a well-known regulator of MT plus-end growth. Here, its localisation suggests that it could stabilise MTs by binding to regions that have been bent by mechanical forces. This possibility could be tested directly by removing the two alleles coding for CLASP2 with CRISPR-Cas9 (see Methods). The absence of CLASP2 was then tested by Western blotting (Figure 4A) and by immunostaining in fixed cells (Figure 4B). Surprisingly, we detected no impact of the absence of CLASP2 on the total amount of MTs or on the amount of NZ-resistant MTs in SCC-untreated cytoplasts (Figure 4C), suggesting that CLASP1 may have compensated for the loss of CLASP2 (*45*). However, the absence of CLASP2 had a striking effect in SCC-treated cytoplasts, in which it completely abolished the mechano-stabilisation of MTs (Figure 4D). The specificity of this effect was tested by overexpressing CLASP2-GFP in CLASP2-KO cells, which was sufficient almost to double the amount of NZ-resistant MTs in SCC-treated cytoplasts (Figure 4E). We have thus demonstrated that CLASP2 has a direct and specific effect on the stabilisation of MTs in response to mechanical forces.

**Figure 4.**
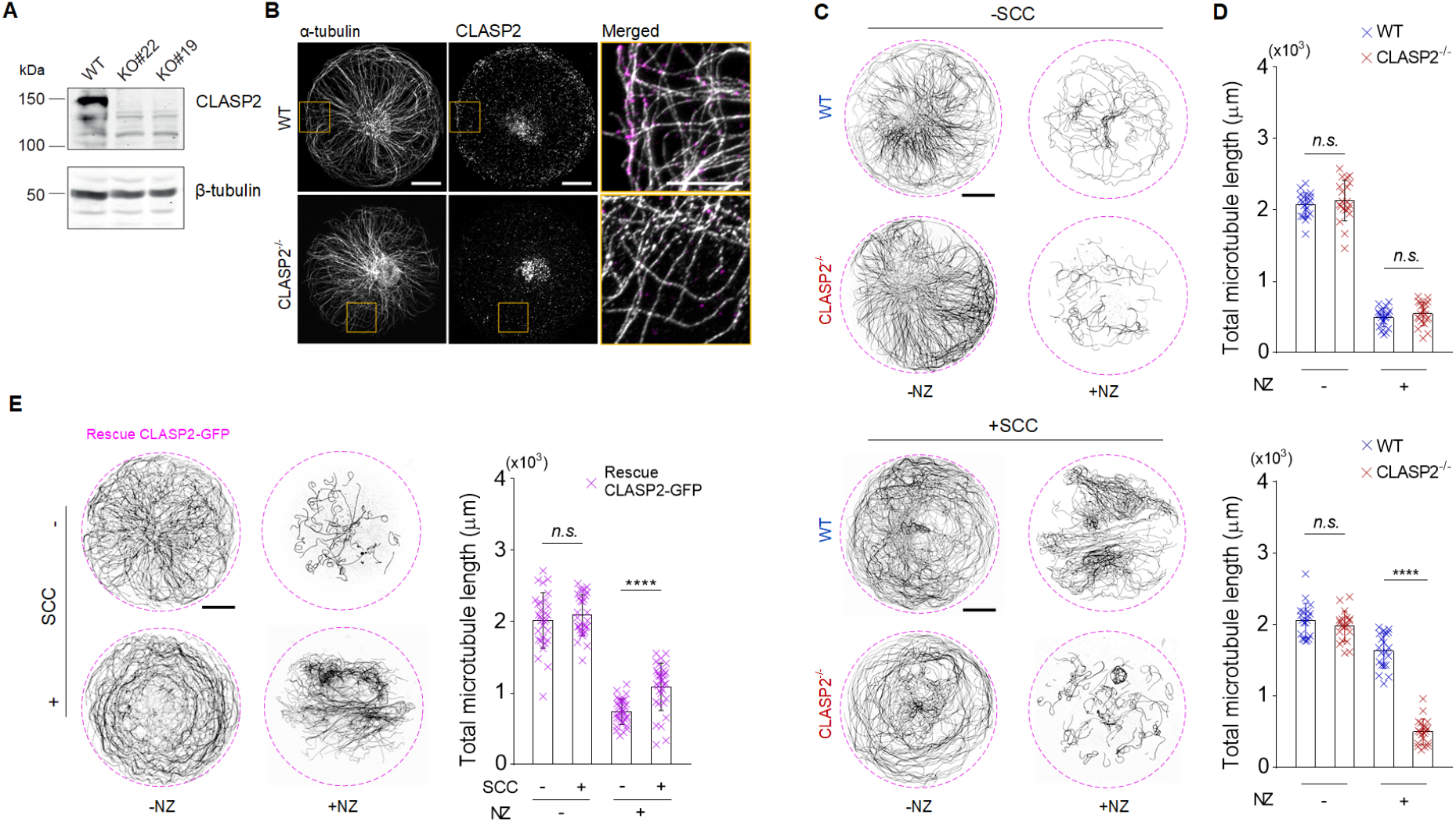
CLASP2 supports microtubule mechano-stabilisation. (A) CLASP2 levels alleles of CLASP2 (two alleles of CLASP2 were removed using CRISPR-Cas9, as KO#22 and KO#19, wild type (WT) as control) in RPE1 cells and measured by immunoblotting. (B) IF images of REP1 WT and CLASP2 knockout (CLASP2^−/-^) cells stained for α-tubulin and CLASP2. (C) Inverted IF images of RPE1 WT and CLASP2^−/-^ cytoplasts in four conditions. (D) Scatter plot of total microtubule length by using SOAX segmentation. (n =20 cytoplasts, three independent experiments). Data are shown as mean ± s.d. Asterisks indicate significance values from the Mann-Whitney test; \*\*\*\**p* < 10^−4^, for *n.s.* (not significant) *p* > 0.1. (E) Inverted IF images of RPE1 cytoplasts expressed by GFP-CLASP2 stained for α-tubulin and the scatter plot of total microtubule length in cytoplasts. (for −SCC/−NZ and +SCC/−NZ conditions: n =30 cytoplasts, for −SCC/+NZ and +SCC/+NZ conditions: n =32 cytoplasts, three independent experiments). Data are represented as mean ± s.d. Asterisks indicate significance values from the Mann-Whitney test; \*\*\*\**p* < 10^−4^, for *n.s.* (not significant) *p* > 0.1. Scale bar, 5 μm (zoomed-in of B) or 10 μm (B, C and E).

In this study, we demonstrated that MTs are mechano-responsive in living cells. We found that cycles of stretching and compression can tear up the actin network and stabilise MTs. Interestingly, our results show that MT stabilisation depends on CLASP2, which relocates from the tip to the shaft of MTs that have been bent by compressive forces. Beyond its role in suppressing catastrophes and promoting rescues at the ends of MTs, CLASP2 is known to repair, protect and stabilise MT shafts when the lattice is physically damaged *in vitro* (*43*). The high curvature we observed suggests that the stress in the lattice exceeded the elastic limit of inter-dimer bonds (*46*) and that the lattice was damaged (*47*), further triggering the local recruitment of CLASP2 to the broken protofilaments (*43*). Interestingly, this re-localisation process is reminiscent of the reinforcement of the actin network by the re-localisation of zyxins from the ends to the damaged part of contractile bundles (*48*–*50*). Overall, these results show that MTs are not rigid rods with a dynamic biochemical exchange at the ends only but that their entire shaft is active (*51*), responding to mechanical forces by deforming and, consequently, recruiting biochemical signals locally.

Stable MTs are present in beating adult cardiomyocytes, where they undergo compression. MT stability and cardiomyocyte contractility are intimately linked (*52*). Pressure overload in the heart has long been known to increase the stability of MTs in cardiomyocytes (*53*, *54*), but the mechanism underlying this regulation is still unclear (*52*). Our results show that the SCCs, such as those associated with a heartbeat, are sufficient to stabilise MT. In turn, stable MTs resist cardiomyocyte contraction (*20*), so the over-stabilisation of MTs is a source of cardiovascular disorders (*55*). Tubulin post-translational modifications, which occur on long-lived MTs, have been shown to impact cardiomyocyte contractility (*20*), and interfering with these modifications of stable MTs is currently a promising strategy to reduce the resistance of stable MTs to cardiomyocyte contraction (*56*). Thus, our identification of CLASP2 as a master regulator of MT mechano-stabilisation may open up new strategies for treatment of cardiomyopathies.

## Acknowledgements

This work was supported by the European Research Council (Consolidator Grant 771599 (ICEBERG) to MT and Advanced Grant 741773 (AAA) to LB), by the Bettencourt-Schueller foundation, the Emergence program of the Ville de Paris and the Schlumberger foundation for education and research. This project was also supported by the MuLife imaging facility, which is funded by GRAL, a program from the Chemistry Biology Health Graduate School of University Grenoble Alpes (ANR-17-EURE-0003). The work of DMR and DV was supported by a grant from the National Institute of Health (R35GM136372). AA was supported by the Netherlands Organisation for Scientific Research (NWO) ECHO Grant 711.018.004. GG was supported by the INCA (AAP PLBIO N°2020-109). MD was supported by the Fondation pour la Recherche Médicale (SPF201809007121)

## Materials and Methods

### Preparation of micro-patterned PDMS membranes

The micro-patterning of cells on silicon membrane was performed according to a deep-UV method previously described in the literature (*57*). Briefly, PDMS substrate (PF-40-X4; GelPak) was firstly washed by 75% ethanol for 5 min on a rotator at 30 oscillations min^−1^ and then incubated with 0.5 mg ml^−1^ poly(L-lysine)-graft-poly(ethylene glycol) (PLL-g-PEG) in 10 mM HEPES buffer, pH 8.6, for 3 h at room temperature. After drying, the modified PDMS samples were stored at 4 °C for at least 1 h. Next, the PLL-g-PEG side of the PDMS substrate was placed on a synthetic quartz photomask bearing a disk shape with 50 μm in diameter. The PLL-g-PEG layer of silicone membrane was exposed to deep-UV (λ=190nm) through the none-chromed windows of the photomask using UVO cleaner (Model no. 342A-220; Jelight), at a distance of 5 cm from the UV lamp with a power of 6 mW cm^−2^, for 5 min. The PDMS membrane was then peeled gently off the mask and incubated with 10 μg ml^−1^ fibronectin from bovine plasma (no. F1141; Sigma) in 10 mM HEPES buffer, pH 8.6, for 20 min at room temperature. For micropattern imaging, PDMS substrate was incubated with a mix of 10 μg ml^−^ 1 fibronectin and 10 μg ml^−1^ fibrinogen–Alexa-Fluor-546 conjugate (no. F13192; Invitrogen) in 10 mM HEPES buffer, pH 8.6, for 20 min at room temperature. Next, the PDMS substrate was rinsed three times with PBS. The micropatterned samples were stored overnight in PBS at 4 °C before plating cells.

### Stretch and compression cycles (SCC)

To enable uniaxial stretching and compression cycles to the micro-patterned cells, we developed an automatic stretcher combined with either a piezoelectric motor (M-663; 18 mm; Linear Encoder; 0.1 μm resolution; PI) or a mechanical motor (M-112.1DG, 25 mm; Linear Encoder; 0.1 μm resolution; PI). Briefly, the silicone grease was plated on top of the micropatterned PDMS membrane (35 mm x 15 mm) to generate a compartment for the culture medium. Then, the PDMS membrane was mounted on the stretcher consisting of a fixed (holding) arm and a mobile (stretching) arm connected to the mechanical motor. Cyclic uniaxial stretches with variable strain (10% to 60%) and frequency (0.1 or 1 Hz) were performed under different conditions. In order to make the SCC assay compatible with confocal microscopy, we used a micromechanical device (*32*). To allow simultaneous stretching of the membrane and to ensure planarity during deformation, we deposited an ultra-thin PDMS sheet (10 μm; Sylgard 184; Samaro DE9330) on a lubricated glass coverslip. To handle such a thin PDMS substrate and avoid mechanical distortion, we enhanced its mechanical stability by adding a thicker (40 μm) elastomeric frame (PF film X0; Gel-Pak) on top of the thin PDMS layer, keeping the size of the observation chamber as small as possible (3 mm × 3 mm; 9 mm^2^). To allow sliding between the PDMS membrane and the glass coverslip, low-viscous glycerol (glycerol for fluorescence microscopy; CAS 56-81-5; Merck) was spun on a plasma-cleaned glass coverslip and then sandwiched with the plasma-cleaned PDMS sheet, creating a glycerol layer of ~0.7 μm thickness. Next, the glass–PDMS sandwich was assembled with a 3D-printed micro-device. The micromechanical device was connected to a piezoelectric motor (M-663; PI) positioned on opposite sides of the observation chamber on the PDMS frame. The observation chamber or the whole microchip could be filled with culture or observation medium.

### Mechanical characterization of the SCC device

We tested numerically and experimentally the mechanical properties of the SCC device. As one of the intrinsic characteristics of uniaxial stretching is that the substrate is stretched along the deformation axis and can be compressed perpendicularly to it, which can be amplified by boundary effects, we wanted to check whether our device allows obtaining a controlled and homogeneous deformation over the observation region. In a first step, we simulated the surface deformation fields for a defined percentage of stretch using a finite element simulation software (COMSOL Multiphysics 4.0a; COMSOL Inc.). The boundary conditions were consistent with the dimensions of the stretching device, and the PDMS substrate was treated as an isotropic, neo-Hookian material with an elastic modulus derived from previous literature (*58*). Our simulations showed that the surface strain fields in the central observation chamber should be remarkably uniform and uniaxial, up to 20% stretch. To experimentally characterise the surface strain fields of the stretched PDMS substrate, mesh grids were marked on the sample. The resulting deformation map is also homogeneous and corresponds well with the simulation data.

### Enucleation of cells

RINZL plastic cover-slides were treated with 1 μg ml^−1^ fibronectin and 12 μg ml^−1^ Collagen I Rat Protein, Tail (no. A1048301, Gibco) for 1hr. The cells were then placed on plastic slides (71890-01, Delta Microscopies) and incubated overnight to achieve an 80% confluence. Next, two plastic slides were placed in a 50 ml tube (no.357007; Beckman) resistant to high-speed centrifugation in complete medium supplemented with Cytochalasin D (no.C8273; Sigma) at 3 μg ml^−1^ for 30 min at 37 °C, then centrifuged at 15,000 g for 1 hr at 37 °C. Enucleated cells were then washed twice with pre-warm DMEM/F12 and incubated for 30 min at 37 °C before being seeded on micro-patterned PDMS substrates.

### Cell culture and seeding

Human telomerase-immortalised RPE1 cells (ATCC), wild-type RPE1 cells, CLASP2 knockout cells, and GFP-EB1 expressing RPE1 cells were cultured in Dulbecco’s Modified Eagle Medium Nutrient Mixture F-12 (no.31331-093; GIBCO) supplemented with 10% fetal bovine serum and 1% penicillin and streptomycin in a humidified incubator at 37 °C and 5% CO_2_. Cells were detached with TrypLE (no.12605036, GIBCO), centrifuged and resuspended in complete medium at 20,000 cells ml^−1^. Cells were plated on the patterned PDMS substrate and allowed to spread for 1 h before washing the unattached cells with a pre-warmed complete medium. The cells were incubated with a fresh culture medium for at least an additional hour at 37 °C to promote correct spreading and polarisation before further treatments. The same culture and plating procedure were applied to the cytoplasts.

### Drug treatment

Microtubules were disassembled by incubating cells/cytoplasts with 2 μM Nocodazole (no. M1404; Sigma) for 15 min (for cells) or 10 μM for 30 min (for cytoplasts) until fixation. Actin filaments were disassembled by incubating cells with 0.1 Cytochalasin D (no.C8273; Sigma) for 20 min. As a control, 0.1 % DMSO (the solvent for either nocodazole or Cytochalasin D) was added to cultures.

### Antibodies, immunofluorescence and immunoblotting

We used rabbit monoclonal antibodies against α-tubulin (no. ab52866, Abcam), monoclonal mouse antibodies against β-tubulin (no. T4026; Sigma), monoclonal mouse antibodies against acetylated tubulin (no. MABT868; Sigma), polyclonal rabbit antibodies against detyrosinated tubulin (*59*), monoclonal mouse antibodies against EB1 (no. 610534; BD Biosciences) and monoclonal rat monoclonal antibodies against CLASP2 (no. KT68; Absea). Primary antibodies were diluted as following for immunofluorescence: α-tubulin (1:500), β-tubulin (1:500), acetylated-tubulin (1:5000), dTyr-tubulin (1:1000), EB1 (1:2000) and CLASP2 (1:500). Secondary antibodies were diluted as following for immunofluorescence: Alexa Fluor 555 goat anti-rat (1:500, no. A21434; Invitrogen), Alexa Fluor 488 donkey anti-mouse (1:500, no. A21202; Invitrogen), Alexa Fluor 488 goat anti-rabbit (1:500, no. A11008; Invitrogen) and Alexa Fluor 546 goat anti-rabbit (1:500, no. A11010, Invitrogen). Primary antibodies were diluted for immunoblotting: β-tubulin (1:5000) and CLASP2 (1:5000). Secondary antibodies were diluted for immunoblotting: IRDye 680LT goat anti-rat and IRDye 800 CW goat anti-mouse (1:5000, Li-Cor Biosciences).

For MT staining, cells were pre-permeabilised in 0.5% Triton X-100 in cytoskeleton buffer for 15 s and then fixed in 0.5% glutaraldehyde (no.00216-30; Polysciences) in cytoskeleton buffer with 0.5% Triton X-100 and 10% sucrose for 15 min at room temperature. Cells were then washed three times with PBS-Tween 0.1% and incubated in a quenching agent of 1mg ml^−1^ sodium borohydride for 10 min at room temperature. For EB1 or CLASP2 immunostaining, cells were fixed in cold methanol at −20 °C for 2 min. For all conditions, after fixation, the cells were washed with PBS-Tween 0.1% and then blocked with 3% bovine serum albumin (BSA) for 1 h. The cells were incubated with appropriate dilutions of primary antibodies in PBS containing 3% BSA and 0.1% Tween overnight at 4 °C in a humid chamber. After washing three times with PBS-tween 0.1%, the PDMS substrates were then incubated with appropriate dilutions of secondary antibodies diluted in PBS containing 3% BSA and 0.1% for 1 h at room temperature in a humid chamber. After washing three times with PBS-Tween 0.1%, coverslips were then mounted onto slides using Prolong Gold antifade reagent containing DAPI for nuclei staining (no. P36935; Invitrogen). The same immunostaining procedure was applied for cytoplasts on the PDMS substrate.

### Generation of CLASP2-knockout cell line using CRISPR/Cas9

20-nucleotide guide sequence targeting exon encoding part of TOG2 domain of CLASP2 was designed using the CRISPR design tool (http://crispr.mit.edu) for the following target sequence: 5’-AGCTAAGGATCTTAGATCCCAGG-3’ Guide sequence was cloned using BbsI restriction sites of pSpCas9(BB)-2A-Puro (PX459), purchased from Addgene (*60*). RPE1 wild type cells were transfected with PX459 plasmid containing the corresponding guide sequence for 24 hours using FuGENE 6 (Promega). Following 72 hours of 4 μM Puromycin selection, cells were recovered and isolated for clone selection. Monoclonal CLASP2-knockout cell lines were characterized and validated by Western blotting and genomic-DNA PCR sequencing using the following primer set: Forward primer: 5’-AGTTTACATTTCTCCGTCGTGC-3’ Reverse primer: 5’-ATATGCAACAACACTGCTTAGG-3’ Sequencing results confirmed generation of a STOP codon in the open reading frame due to a frameshift mutation created by the deletion of 111th bp in exon 11. This led to the translation of only 392 aa residues of CLASP2 (UniProtKB - E7ERI8).

### Cell transfection

GFP-CLASP2 plasmids are cloned from *H. sapiens* and constructed in pEGFP-C1 by linking the 5’ portion of the human truncated EST clone 7k43h10.x1 (IMAGE:3478506) to the nucleotides 194–5614 of the KIAA0627 cDNA (*61*). GFP-CLASP2 were transfected into cells using an X-tremeGENE kit (*62*) as per the manufacturer’s instructions 48 h prior to the experiment. The transfected cells were then put to the enucleation procedure.

### Imaging

Images of the different immunostainings were acquired on a Zeiss LSM880 or a Zeiss LSM900 confocal microscopes (Axio Observer) using either a 63x magnification objective (Plan-Apochromat 63X/1.4 oil) or a 20x magnification objective (Plan Apochromat 20x/0.8). Only cells that were well spread out on disk micropatterns were selected for imaging.

Image acquisition for time-lapse of GFP-EB1 was performed on an inverted microscope (DMi6000; Leica) equipped with a spinning disk unit CSU-X1 (Yokogawa Electric Corporation; Japan) through oil-immersion objective (Leica, HC PL APO 63x/1.40 oil) objective every 1 s during 2 min for each time-lapse. The set-up was equipped with a live cell chamber, and the temperature was constantly kept at 37 °C. EB1 comets were excited with a 491 nm laser line, and emission was observed with a standard GFP filter. The microscope was monitored with MetaMorph software (Universal Imaging). To register fluorescence images and measure PDMS deformation, we adsorbed 0.1 μm fluorescent beads (TetraSpeck Microspheres; Thermo Fisher Scientific) on the SCC chamber that were imaged during the entirety of the microscope acquisitions.

### Image analysis

#### Microtubule segmentation

Microtubules in cells were segmented using SOAX software (v. 3.7) (*44*). Briefly, the underlying SOAX method uses multiple stretching open active contours that are automatically initialised at image intensity ridges and then stretched along the centrelines of filaments in the network. Microtubule density was determined by the ratio of the total number of microtubules in individual cells segmented by SOAX and cell spreading area. The total length of microtubules in the cells was calculated by summing the length of SOAX-segmented microtubules. The same segmentation analysis was performed for the cytoplasts. For the quantification of small microtubules in cytoplasts, the raw data were converted to 8-bit grayscale images and then analysed with an Image Pro-Plus cluster plug-in (v. 6.0; Media Cybernetics).

#### Quantification of the cortical microtubule deformation

To quantify the quasi-periodic deformation of cortical microtubules upon SCC, we analysed their spatial wavelength similarly to the previously described method (*63*). Briefly, we obtained the orientation of the microtubules by evaluating the gradient structure tensor as implemented in the OrientationJ (*64*) plug-in for FiJi (1.52v) (*65*). Using a custom MATLAB (v. 2018a, MathWorks Inc.) script, with the aid of MATLAB-FiJi interface MIJ (*66*), we extracted four annulus sectors corresponding to the cortical regions (angular length of each sector was 0.6 rad and the side length 0.6 μm) from the orientation field. These sectors were paired: one pair corresponding to the cortical part, which did not experience circumferential strain, and the other pair, which underwent maximum circumferential strain during SCC. In these pairs, using Fast Fourier Transform, we calculated the mean spectrum of orientation along the circumferential direction, which was corrected for discontinuities arising from the modulo operation of the angle assignment. We measured the wavelengths corresponding to the most prominent non-zero mode of the obtained spectra. These wavelengths are, due to the linearity of the Fourier transform, equivalent to the principal wavelengths of the microtubule deformation.

#### Geometric transformation of images

To facilitate analysis and visualise the microtubule orientation, we applied a geometric transformation as shown in Fig. 2A. This conformal mapping transforms a disk into a rectangle in a way that (i) radial lines become parallel line segments, and (ii) circles concentric with the disk become line segments as well. The two types of resulting line segments, (i) and (ii), are perpendicular as a result of conformity. The centre of the disc, which can be treated as a circle with a zero radius, maps on a line segment similarly to other concentric circles. To apply this transformation to the centred images of cells on micropatterns, we used a numerical method written in MATLAB. The fluorescence intensity profiles along the circular path concentric with the cell were taken sequentially with increasing radius. Each profile was resampled using a linear anti-aliasing FIR filter to obtain profiles of identical lengths. The resulting profiles were arranged in a matrix and visualised.

#### EB1 Comet tracking and Analysis

EB1 comets moving speed and duration time were quantified using a TrackMate plug-in for Fiji (*67*). To build average intensity distribution at the microtubule growing tip, we generated intensity profiles of 6 pixels thick line (200 nm) of 4-5 μm length along the axis of microtubule at the MT tip using Fiji. The fluorescent profile was then normalised by the maximum intensity value.

#### Quantification of the orientation variance of microtubules

To visualise and quantify the variation of microtubule orientations from all cytoplasts imaged at a particular experimental condition, we used the variance of microtubule orientation (*68*) of the mean projection of microtubules. The orientation vector field of microtubules was obtained using OrientationJ as described above. A sliding window was then applied to this field in MATLAB. Within each window position, we calculated the mean resultant length of the orientation unit vectors 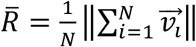, and the circular orientation variance as 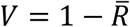. To visualise the result, the matrix of orientation variances was resized using the bicubic interpolation method, and the parts of the field outside of the cell were masked by thresholding the original average image.

#### Co-localisation between MT and CLASP2

To determine the coupling state of CLASP2, we used super-resolution reconstruction of the fluorescence images. Microtubules were segmented from the 488 nm channel of fluorescence images using SOAX (v. 3.7) (*44*). This method extracts centrelines of microtubules which we interpret as the likeliest estimate of microtubules’ axes. The CLASP2 was segmented from the 555 nm fluorescence channel using the FIESTA (v. 1.05) particle tracking software (*69*), which estimates the positions of the molecules from the 2D Gaussian approximation for the point spread function. Next, we deleted microtubule and CLASP2 detections that fall within the nuclear region, which we identified by thresholding the fluorescence image in the DAPI channel. The coordinates of the remaining microtubules and CLASP2, which we interpret as localised within the cytoplasm, were compared in MATLAB to determine their mutual distances. For each cytoplasmic CLASP2 detection, we calculated the shortest distance to the nearest microtubule axis. We considered CLASP2 as microtubule-associated if the particle was confined within the coupling radius from an axis of a microtubule. This radius was estimated from the physical size of the construct of CLASP2 (7.5 nm, (*70*)), primary and secondary antibodies (20.1 nm) (*71*) and the microtubule geometry (radius 12.5 nm) (*72*) as 40.1 nm. The CLASP2 was regarded as uncoupled otherwise.

### Statistical analysis

Statistical analyses were performed using Graphpad Prism 6 software (GraphPad Inc.) and R studio v 1.4. Respective numbers of data points, n, are shown in the figure legends. The indicated P values were obtained using a two-tailed unpaired Mann Whitney test or Kruskal-Wallis test with Dunn’s multiple comparison test.

---

**Supplementary Figure 1.**
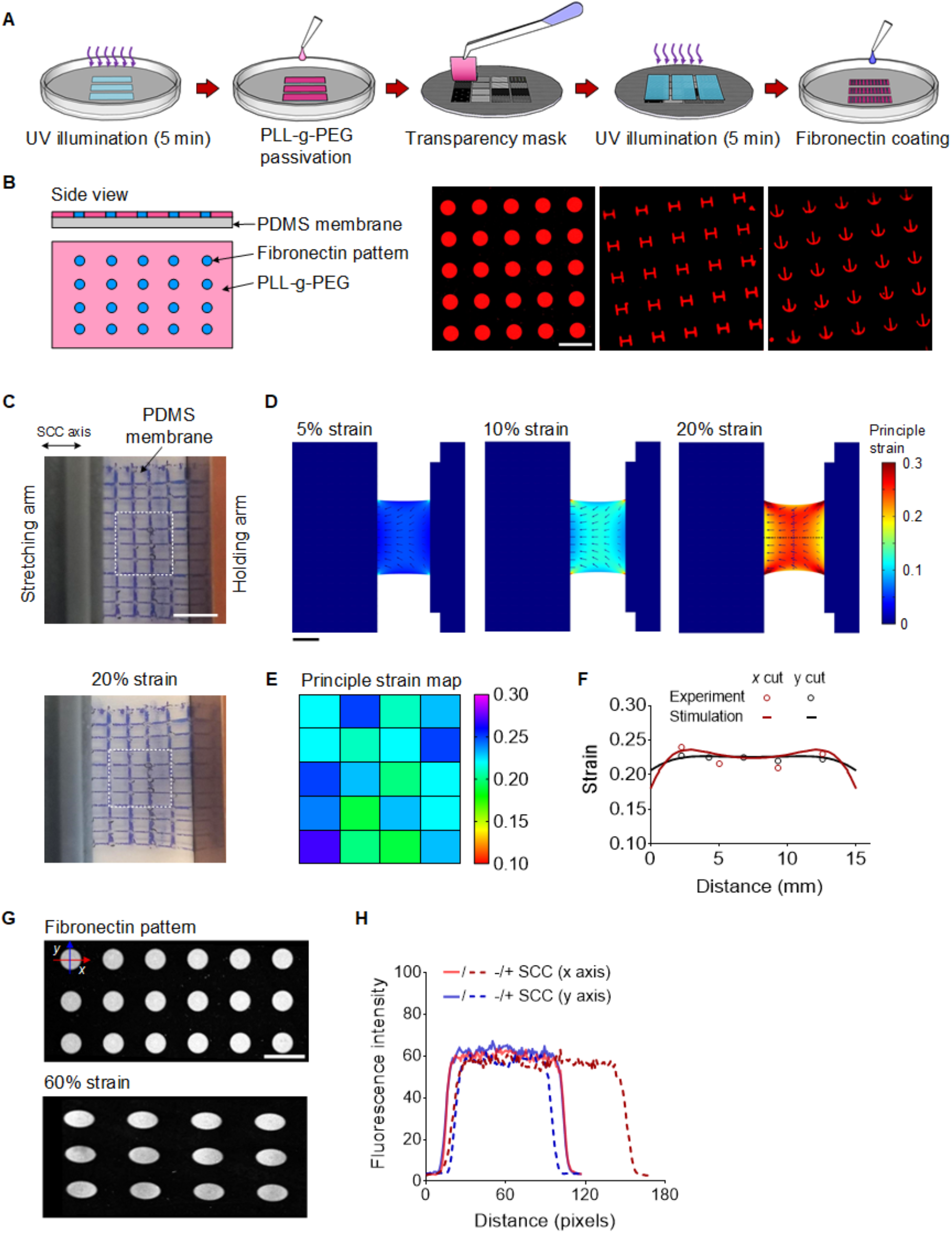
Characterization of SCC assay combined with micropatterning. (A) Illustration of micropatterning process on PDMS membrane. (B) Illustration of surface properties of a PDMS membrane with fibronectin micropatterns (left, labelled by fibrinogen– Cy3) and IF images of L-, H- and crossbow-shaped micropatterns of fibronectin. (C) Images of grid-labelled PDMS membrane held by a stretcher composed of a holding and stretching arm (top) and applied up to 20% stretch. (D) Numerical simulation by using COMSOL Multiphysics of 5% (left), 10 % (middle) and 20 % (left) stretches for a PDMS membrane (150 μm in thicknesses). The first principal strain is colour coded from 0 to 30 %, and arrows display displacement fields (XY plane). (E) Colour-coded map of principle strain obtained by calculating the displacement variations of grids on PDMS membrane before and after 20% stretch. (averaging from 5 samples of two independent experiments) (F) Line profile of the simulated first principle strain along the principal axes of the stretching PDMS membrane, which are in good fitting with the experimental data (spots). In both axes, the strain was in large parts nearly constant (for 5 mm < x,y < 10 mm). (G) IF images of fibronectin micropatterns (labelled by fibrinogen–Cy3) before and after 60% stretch. (H) Line-scan profiles of fluorescence intensity along the *x* and *y* axes of micropatterns before and after 60% stretch. (averaging from 14 patterns of three independent experiments). Scale bar, 100 μm (B and G) or 5 mm (C and D).

**Supplementary Figure 2.**
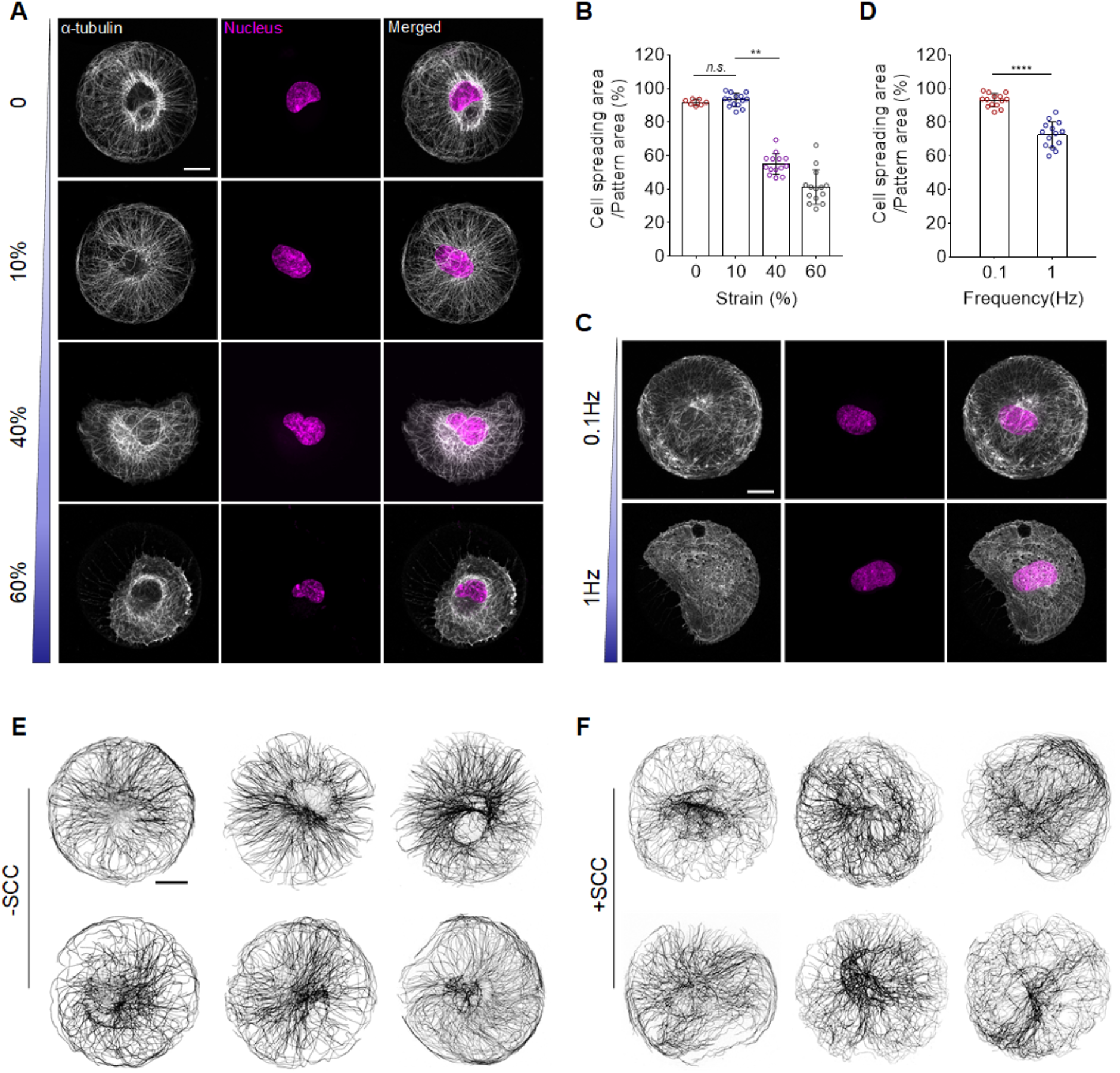
Cell resistance to various strains and rates of SCC. (A) IF images of RPE1 cells applied with 0, 10%, 40% and 60% SCC at 0.1 Hz, stained for α-tubulin (grey) and DNA (magenta). (B) Scatter plot of the ratio of cell spreading area and micropattern area with varying SCC. Data are shown as mean ± s.d. Asterisks indicate significance values from Kruskal-Wallis test with Dunn’s multiple comparison test; \*\**p* < 10^−2^, for *n.s.* (not significant) *p* > 0.1. (C) IF images of RPE1 cells applied with 10% SCC at 0.1 and 1 Hz, respectively, stained for α-tubulin (grey) and DNA (magenta). (D) Scatter plot of the ratio of cell spreading area and micropattern area with varying SCC frequency. Data are shown as mean ± s.d. Asterisks indicate significance values from the Mann-Whitney test; \*\*\*\**p* < 10^−4^. (E-F) Representative IF images of RPE1 cells applied with and without SCC (10% strain at 0.1 Hz), stained for α-tubulin (grey). Scale bar, 10 μm.

**Supplementary Figure 3.**
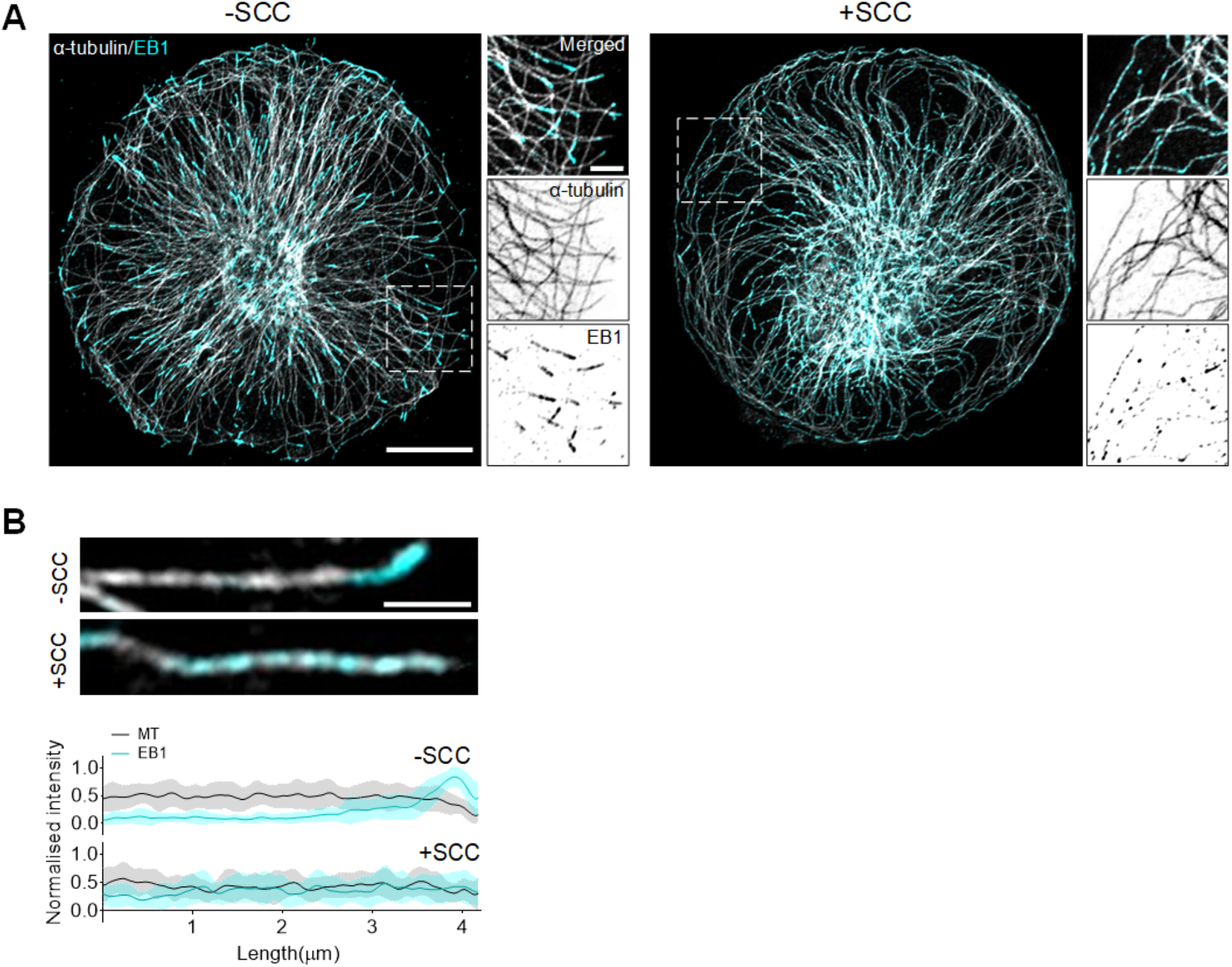
Characterization of EB1 relocation in RPE1 cells in response to SCC. (A) IF images of RPE1 cells without/with SCC, stained for α-tubulin (grey) and EB1 (cyan). Dashed rectangle designates region shown with merged, α-tubulin or EB1 channel isolated. (B)Representative IF images of individual microtubules and EB1 comets in cytoplasts without/with SCC (top) and corresponding line-scan profiles of normalised fluorescence intensity (bottom) (n =20 individual microtubules from 10 cells, three independent experiments). Means are depicted. Scale bar, 10 μm (A), 2 μm (zoomed-in of A) or 1 μm (B).

**Supplementary Figure 4.**
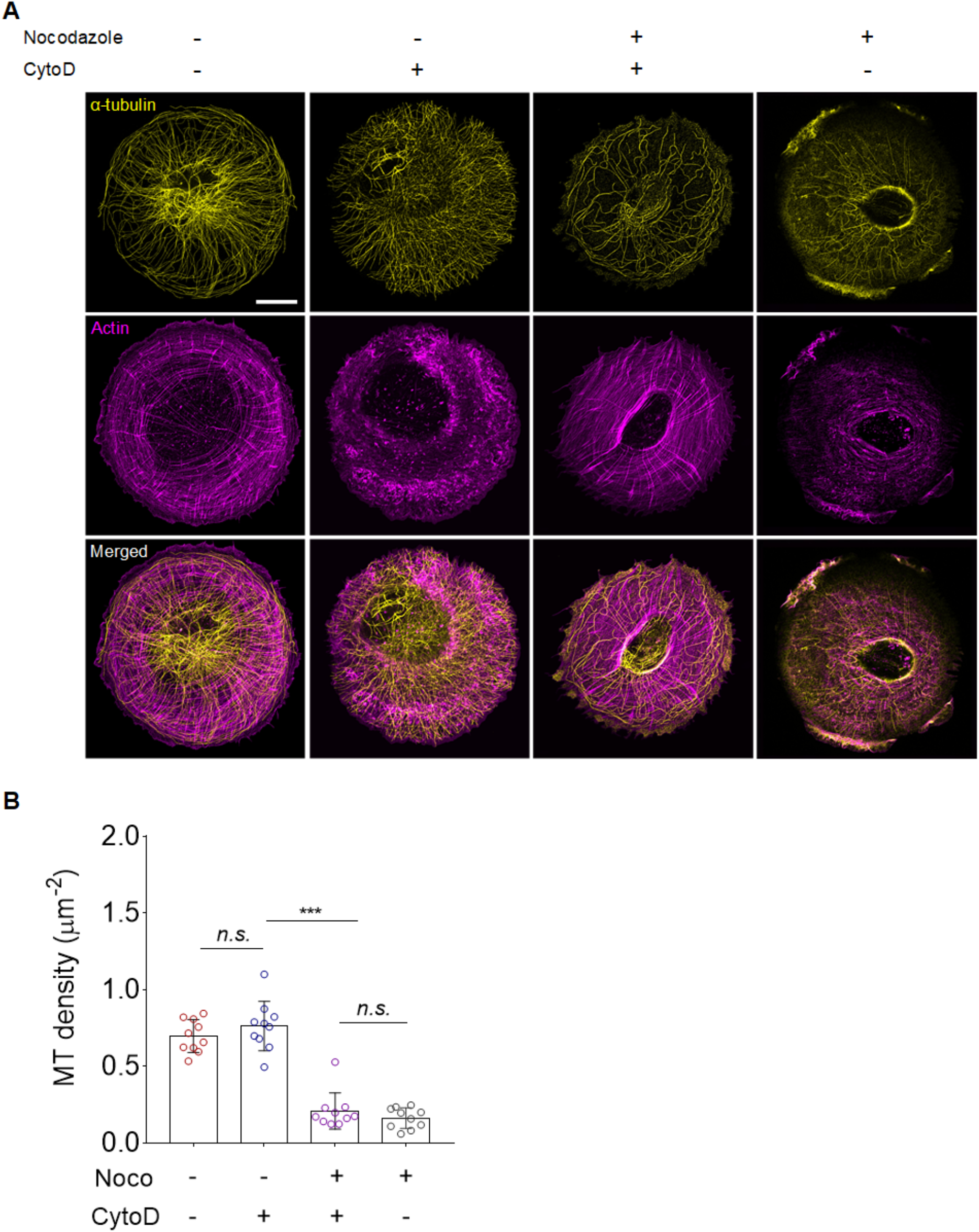
Actin disassembly is not sufficient to stabilise MTs. (A) IF images of RPE1 cells treated with nocodazole and cytochalasin D, stained for α-tubulin (yellow) and actin (magenta). (B) Scatter plot of the ratio of microtubule density of RPE1 cytoplasts in varying conditions. (n =10 cells, three independent experiments). Data are shown as mean ± s.d. Asterisks indicate significance values from Kruskal-Wallis test with Dunn’s multiple comparison test; \*\*\**p* < 10^−3^, for *n.s.* (not significant) *p* > 0.1. Scale bar, 10 μm.

**Supplementary Figure 5.**
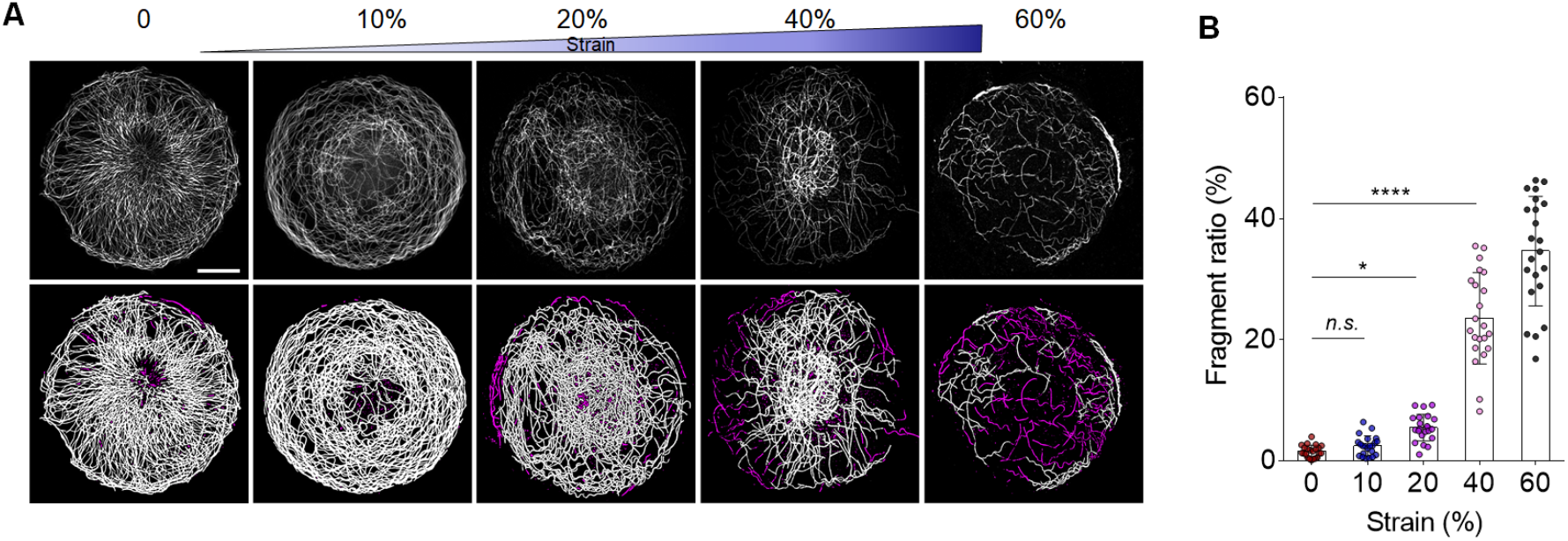
Resistance of the MT network of the enucleated cell (cytoplast) to various rates of SCC. (A) IF images of RPE1 cytoplasts subjected to 0, 10%, 20%, 40% and 60% SCC at 0.1 Hz, stained for α-tubulin (grey). (B) Scatter plot of the fragment ratio of microtubules with varying SCC. (n =22 cytoplasts, three independent experiments). Data are shown as mean ± s.d. Asterisks indicate significance values from Kruskal-Wallis test with Dunn’s multiple comparison test; \*\*\**p* < 10^−3^, \**p* < 0.05, for *n.s.* (not significant) *p* > 0.1. Scale bar, 10 μm.

**Supplementary Figure 6.**
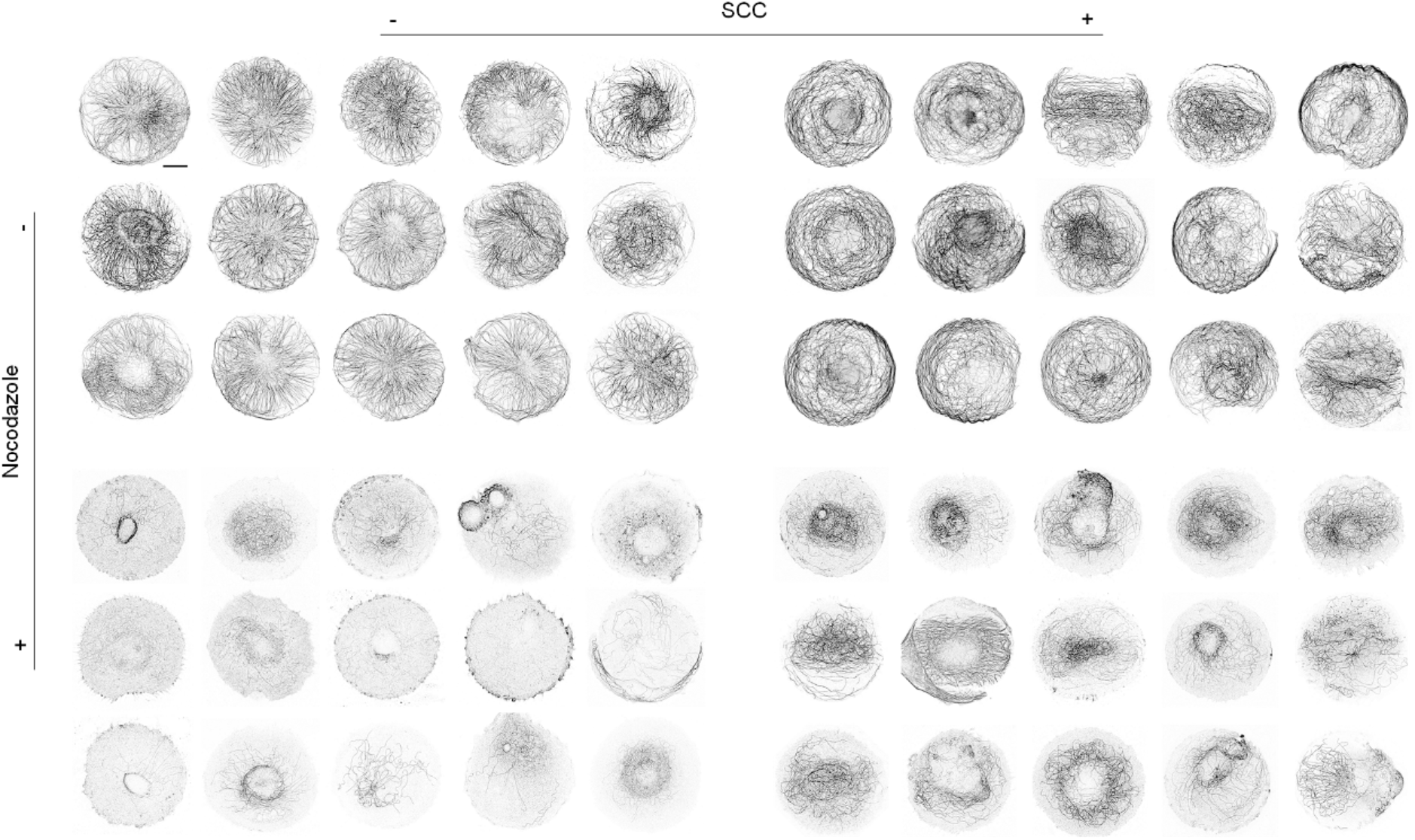
MT resistance to nocodazole in enucleated cells (cytoplast) in response to SCC. (A) Representative inverted IF images of RPE1 cytoplasts stained for α-tubulin in varying conditions. −SCC/+SCC: without/with SCC, −NZ/+NZ: without/with nocodazole treatment. Scale bar, 10 μm.

**Supplementary Figure 7.**
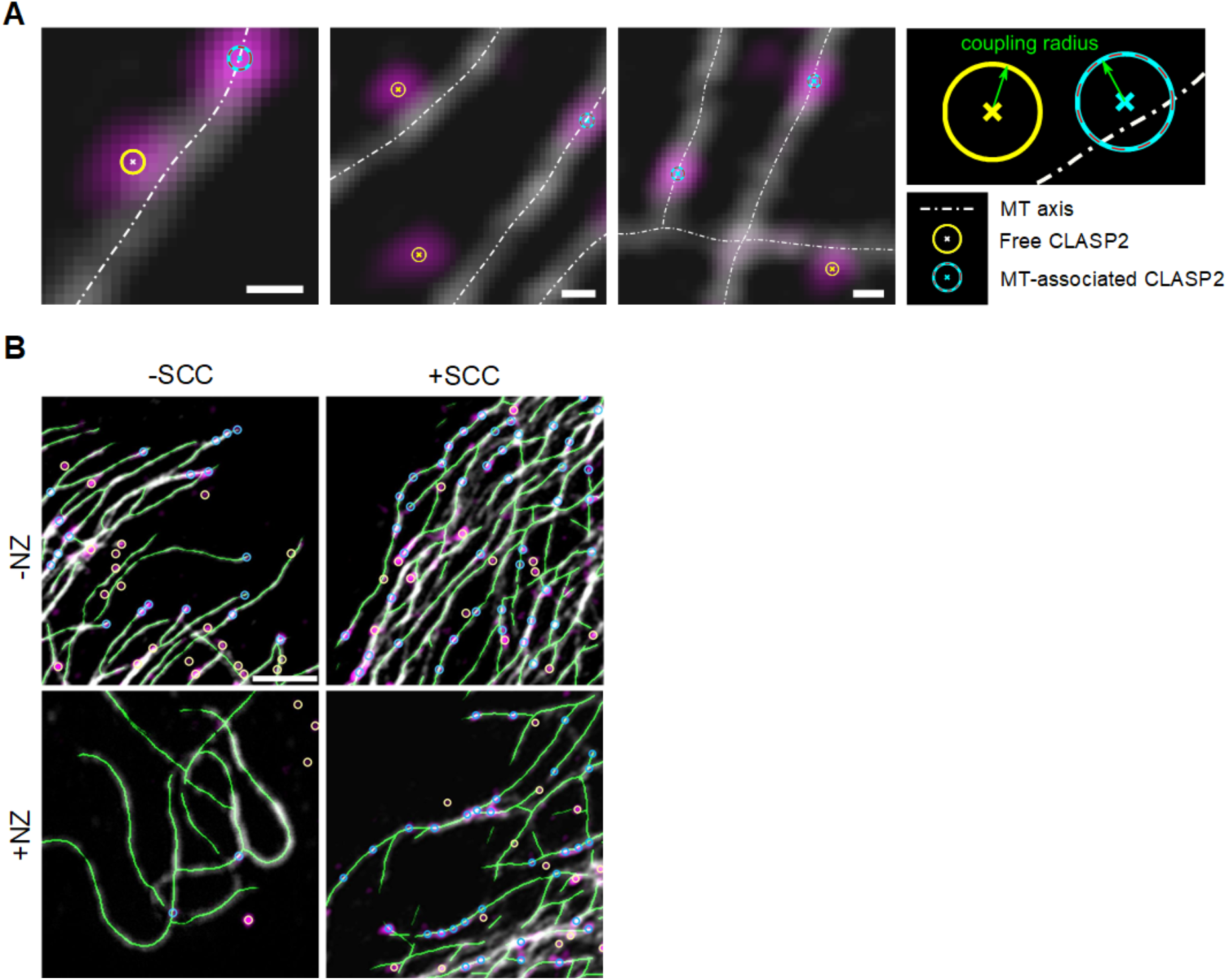
Illustration of the segmentation method to quantify CLASP2 association to MTs. (A) Representative IF images of microtubules in RPE1 cytoplasts stained for α-tubulin (grey) and CLASP2 (magenta). The microtubule-associated (coupled) or free (uncoupled) CLASP2 particles are marked by yellow and blue circles, respectively. The axis of microtubules is depicted as a white dash-dot line. For each detected cytoplasmic CLASP2, its association with the microtubule is determined by comparing the shortest distance to the nearest microtubule axis with the coupling radius. (B) Representative images of detected CLASP2 and microtubules in varying conditions. Scale bar, 200 nm (A) or 2 μm (B).

